# Random Forest approach for the identification of relationships between epigenetic marks and its application to robust assignment of chromatin states

**DOI:** 10.1101/2023.01.12.523636

**Authors:** Leandro Murgas Saavedra, Gianluca Pollastri, Erick Riquelme, Mauricio Sáez, Alberto J.M. Martin

**Affiliations:** Programa de Doctorado en Genómica Integrativa, Vicerrectoría de investigación, Universidad Mayor, La Pirámide 5750, Huechuraba, Chile; Laboratorio de Redes Biológicas, Centro Científico y Tecnológico de Excelencia Ciencia & Vida, Fundación Ciencia & Vida, Facultad de Ingeniería, Arquitectura y Diseño, Universidad San Sebastián, Avenida Zañartu 1482, Santiago, Chile; School of Computer Science, University College Dublin, Belfield, Dublin 4, Ireland; Department of Respiratory Diseases, Facultad de Medicina, Pontificia Universidad Catolica, Santiago, Chile; Centro de Oncologia de Precisión, Facultad de Medicina y Ciencias de la Salud, Universidad Mayor, Santiago, Chile

## Abstract

Structural changes of chromatin modulate access to DNA for all proteins involved in transcription. These changes are linked to variations in epigenetic marks that allow to classify chromatin in different functional states depending on the pattern of these marks. Importantly, alterations in chromatin states are known to be linked with various diseases. For example, there are abnormalities in epigenetic patterns in different types of cancer. For most of these diseases, there is not enough epigenomic data available to accurately determine chromatin states for the cells affected in each of them, mainly due to high costs of performing this type of experiments but also because of lack of a sufficient amount of sample or degradation thereof.

In this work we describe a cascade method based on a random forest algorithm to infer epigenetic marks, and by doing so, to reduce the number of experimentally determined marks required to assign chromatin states. Our approach identified several relationships between patterns of different marks, which strengthens the evidence in favor of a redundant epigenetic code.

## 1 Introduction

The regulation of gene expression is a crucial factor in the development and maintenance of life in all organisms. Gene expression is mainly regulated at the transcription level, controlling the quantity of mRNA produced, and at translation governing the quantity of protein synthesized and the number of products encoded by each gene. Importantly, errors in any of the two levels of gene regulation is often responsible for many diseases.

Structural changes of chromatin modulate access to DNA of all proteins involved in transcription. This process is linked to variations in histone modifications which are also known to be related to an ATP-dependent remodeling complex that causes the dynamic restructuration of nucleosomes [1]. Histone modifications are covalent Post-Translational Modifications (PTMs), usually located in the histone tails, that include methylation, phosphorylation, acetylation, ubiquitylation, and sumoylation [2]. These PTMs are known to affect chromatin structure and can be classified as active or repressive marks based on their effect on transcriptional regulation. For example, active marks, such as acetylation of lysine 9 and di-methylation of histone 3 (H3K9ac and H3K4me2, respectively), are related to chromatin de-condensation and the formation of chromatin loops that allow access to DNA by the transcription machinery [3]. Another well characterized example is the X chromosome inactivation by numerous repressive marks, such as tri-methylation of lysine 27 and di-methylation of lysine 9 of histone 3 (H3K27me3 and H3K9me2 respectively) [4].

The hypothesis of the “histone code” was proposed in the early 2000s [6]. This histone code states that a combination of histone modifications at a specific genomic locus determines the activity of the underlying gene. Subsequently, the histone code was expanded to “epigenetic code” to include other epigenetic marks such as DNA methylation [7, 8]. The epigenetic code is considered conserved in each species, that is, the effect of a pattern of marks is identical for all cells of the same organism, and all individuals of the same species [2]. To give an example, the promoter of the same gene in cells of different tissues can be active or inactive, and in the same way, this promoter can be associated with different patterns of histone modifications. Currently, it is thought that the epigenetic code is redundant, different combinations of histone modifications contain the same message [2]. Additionally, different studies show that existing patterns of histone modifications are also strongly related to the structure of chromatin [6, 9].

Chromatin was traditionally divided into two functional states, one transcriptionally active (euchromatin) and one inactive (heterochromatin) [10]. More recently, newer chromatin annotation approaches have classified chromatin into different states, ranging from two types to several tens, each associated with a different functional condition [11]. For example, a large-scale, integrative genome-wide analysis of 53 chromatin-associated proteins in *Drosophila melanogaster* recognized five chromatin states classified and named by colors [12]. In this classification of five states, the transcriptionally active state is divided into three (green, yellow and red) and the inactive state into two (blue and black). Most classifications of chromatin states are based on different criteria, most of which rely on epigenomic information data for many histone marks and DNA methylation [13, 14].

One of the most widely used tools for to classify chromatin states is ChromHMM [5]. This methodology is based on an unsupervised tool which assigns states to chromatin using epigenetic information in one or more cell types. Using this approach, annotation is carried out employing multivariate Hidden Markov Models based on the absence or presence of epigenetic marks in the genome. Through this methodology, using information from different epigenetic marks in various cell lines, the authors manage to assign 18 states to chromatin, thus further expanding the number of known states [15].

Many diseases have been related to changes in chromatin states associated to misregulation of gene expression [16]. These abnormal variations in gene regulation are caused by the alteration of diverse epigenetic factors, mutations in specific genes, or regulatory regions of chromatin. Among these diseases, it is possible to highlight: most types of cancer, diabetes, and several neurological and cardiovascular complex diseases [17, 18, 19]. For instance, several studies have associated colorectal cancer with alterations in chromatin states identifying abnormalities in DNA methylation patterns and different histone modifications, such as tri-methylation of lysine 4 and 9 of histone 3 (H3K4me3 and H3K9me3) and tri-methylation of lysine 20 of histone 4 (H3K27me3) [20, 21]. Melanoma progression has also been associated with variations in epigenetic patterns, where it has been observed that there is a loss of histone acetylation and di and tri-methylation of lysine 4 of histone 3 (H3K4me2/me3) in regulatory regions, which would be affecting the development of the disease [22]. Importantly, for most of these diseases, there is not enough epigenomic data available to accurately determine chromatin states for the cells affected in each of them.

Machine Learning (ML) has increasingly being applied to the study and diagnosis of diseases. The main reason behind this is its great ability to analyze large amounts of data and find patterns which would be difficult to detect with the naked eye [23]. ML has proven to be indispensable for interpreting large genomic datasets and integrating several types of “omic” data. In this way, ML methods can be used to learn how to recognize the location of Transcription Start Sites (TSSs) in a genome sequence [24], infer gene expression [25] or annotate new Transcription Factor Binding Sites (TFBSs) [26]. ML algorithms can use input data generated by different genomic assays, such as expression data, chromatin accessibility assays, histone modifications, or TFBSs [23]. There are several ML based methodologies that help to solve the problem of scarcity of experimental information. For example, CHROMIMPUTE [27] and AVOCADO [28] which use ML-type strategies to predict PTM enrichment signals from epigenetic information. CHROMIMPUTE relies on regression tree-type algorithms to carry out its imputations based on information on PTMs present in the target sample as well as in others previously trained. Similarly, AVOCADO uses multi-scale deep tensor factorization-oriented techniques which represent epigenetic information in a neural network model that allows predictions of PTMs. It should be noted that both methodologies use a large amount of information to make optimal predictions and in the case of those oriented towards deep learning, they could be difficult to employ by non-expert users. In addition, the information provided by these tools only indicates the presence or absence of the target PTM, without explaining how this prediction was made or what relationships can be found between the different data used, important information to for example understand the epigenetic code.

With the purpose of overcoming the lack of explainability of other approaches, in this study we describe a new method based on the Random Forest (RF) algorithm to predict histone modifications. Notably, due to the inherent properties of this algorithm, our approach indicates the different existing relationships between histone PTMs that are employed to be predict one PTM based on the others. This new tool allows to reduce the number of experiments necessary for the assignment of chromatin states, increasing the speed of the analyses and reducing the costs associated with them.

## 2 MATERIALS AND METHODS

### 2.1 Data

Different types of data were used to generate datasets. Enhancer annotations on the genome were obtained from the database GeneCards [29], and promoters and gene coordinates from ENSEMBL [30], all data in human genome version GRCh38. In the case of PTMs, data were obtained from a study of Fiziev et al. [22] which consists of ChIP-seq experiments for 33 histone PTMs in two biological conditions, the tumorigenic (Tum) and non-tumorigenic (noTum) phenotype of melanocytes (GSE58953) (each condition has two biological replicates, Hmel and Pmel cell lines). The Fastq files of the ChIP-seq experiments were reanalyzed and aligned against the human genome GRCh38 using Bowtie2 [31]. PCR duplicates were removed using SAMTools [32], and BEDgraph files were obtained with the BamCoverage tool from the Deeptools suite [33], using the BAM files for the 33 PTMs with parameters by default, different bin sizes, and normalizing the reads to RPKM. For colorectal cancer data, ChIP-seq experiments of 11 PTMs of the HCT116 cell line available in ENCODE were analyzed with the same pipeline before using them as test set.

### 2.2 Datasets

Once the ChIP-seq experiments were processed, they were used to create datasets to train and test the ML algorithms (Fig 1.A). The genome is segmented in fragments of *n* base pairs (different values of *n* were tested and chosen according to performance). Each fragment is then described by a numerical vector where each element represents numerically the absence or presence epigenetic mark associated with the said fragment. For example, the absence/presence of PTM was indicated by the RPKM value per fragment of reads in the experiment aligned against it. Labels were assigned in numbers between 0 and 1, using the “clipping” technique, in which those fragments with RPKM values less than 2 were divided into 2, and those greater than or equal to 2 were assigned a value of 1. Also, a distance (*d*) upstream and downstream of the fragment to be predicted was used to observe the relevance of each mark in the flanking regions. Finally, each vector is associated with a label representing the characteristic to predict, for example, a specific PTM.

**Figure 1:**
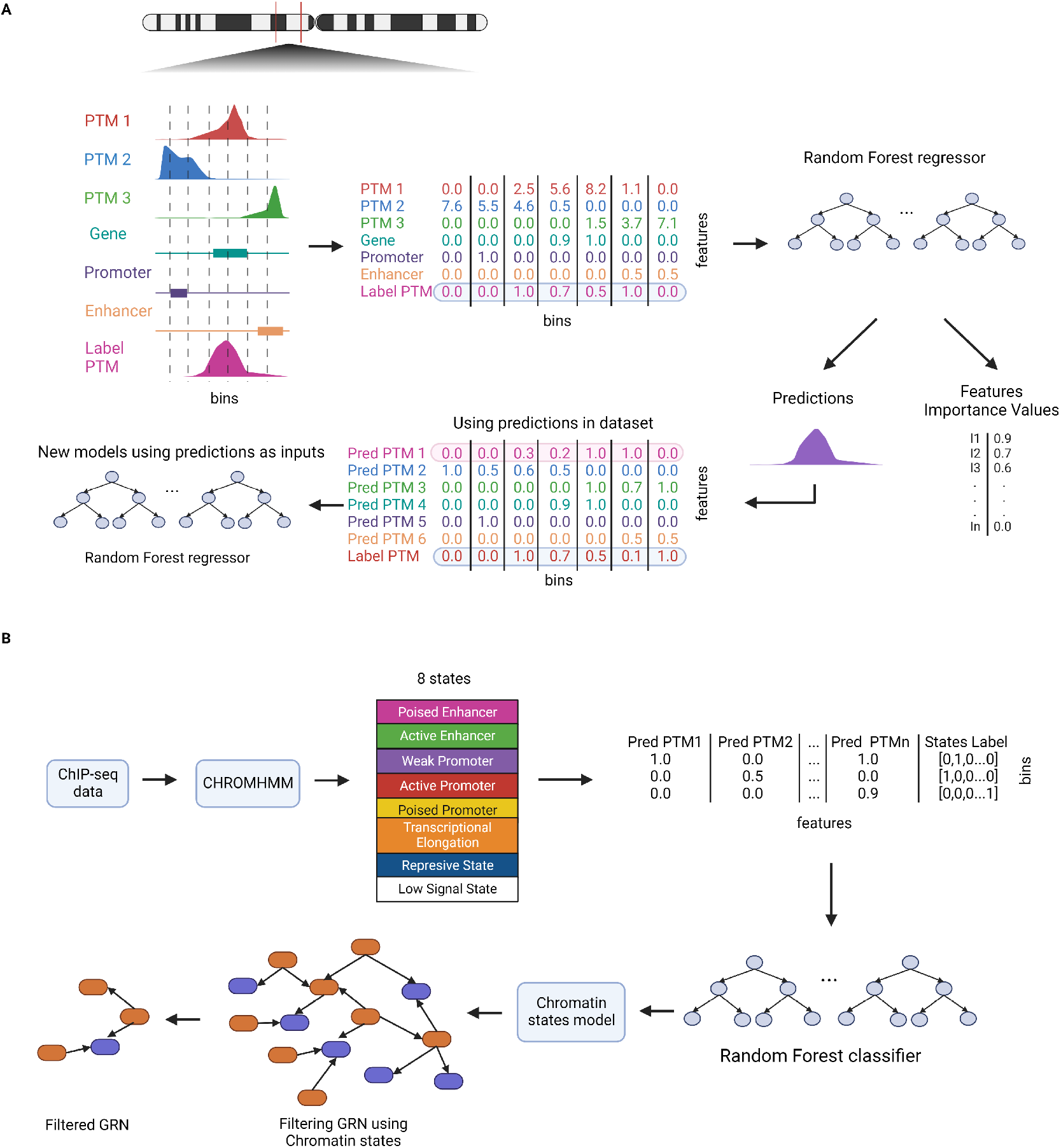
Scheme of the methodology used. (**A**) We first created different datasets for the identification of relationships between PTMs using data from ChIP-seq experiments, enhancers, genes and promoter annotations. These datasets were next employed for the training and testing of RF regression algorithms. For this, the genome was divided into fragments (bins) of size *n* and the presence or absence of the different features was represented quantitatively using RPKMs per fragment in the case of the ChIP-seq experiments, for the other data, the percentage of coverage of the data on the fragment. In this way, it is possible to predict the different PTMs and determine the most important features, managing to identify the different relationships between PTMs. Additionally, the predictions obtained were used to generate new models that are based on a reduced number of experimental data types. (**B**) In a second stage of our approach we employed predicted PTMs for the assignment of chromatin states. First, ChromHMM was used for the assignment of states with experimental information, assigning 8 states to the chromatin. Then this assignment was coded for the training and testing of a RF classifier, which allows classifying the different states based on the predicted data. Finally a reference GRN was filtered according to the chromatin state assignment based on the regulatory nature associated to that state.

### 2.3 Parameters Determination

We employed RF regressors as implemented in Scikit-Learn [34], trained and tested using the 32 PTMs ChIP-seq data as input in addition to Enhancer, Genes, and promoters annotations, to predict each of the PTM one at the time. We performed all the experiments used to determine the RF training parameters using solely chromosome 1 data. This chromosome was chosen because it is the largest one and contains significant enough examples. Different training parameters were tested to define fragment size, distance and the RF depth (see Table 1), using 75% of the dataset to train each of the combination of parameters and the 25% remaining as test set. The performance is evaluated with Pearson’s correlation coefficient between the label of the data used for testing and the predicted values, so values close to one indicate a higher performance in the model.

**Table 1:** Parameters RFs. Parameters used to identify the most adequate way to represent the information, First column indicates the fragment size; Second column the distances adjacent to the central fragment; Third column the depth value used to train the RF algorithm

| Fragment | Distance | Depth tree |
| --- | --- | --- |
| 200 | 2000 | 15 |
| 500 | 5000 | 25 |
| 1000 | 10000 | 50 |
| 2000 | 15000 | Max |

Our first analysis used three PTMs, H3K4me1, H3K4me3, and H3K27ac, with the Pmel-noTum cell line data. The results show Pearson’s correlation values between 0.88 and 0.95 for H3K4me1, 0.82 and 0.93 for H3K4me3, and in the range 0.94 and 0.98 for H3K27ac. In addition, we found that the size of the fragment is the hyperparameter with the most impact, since by varying it the performance of the model changes considerably, unlike with the values of distance or tree depth (See Table 2 for the case of models generated with depth 25, in the case of other depths see Supplementary Table 1). According to these results, we decided to generate models for the 33 available PTMs using a fragment size of 2K bases, distances of 10K bases, and tree depth 25.

**Table 2:**
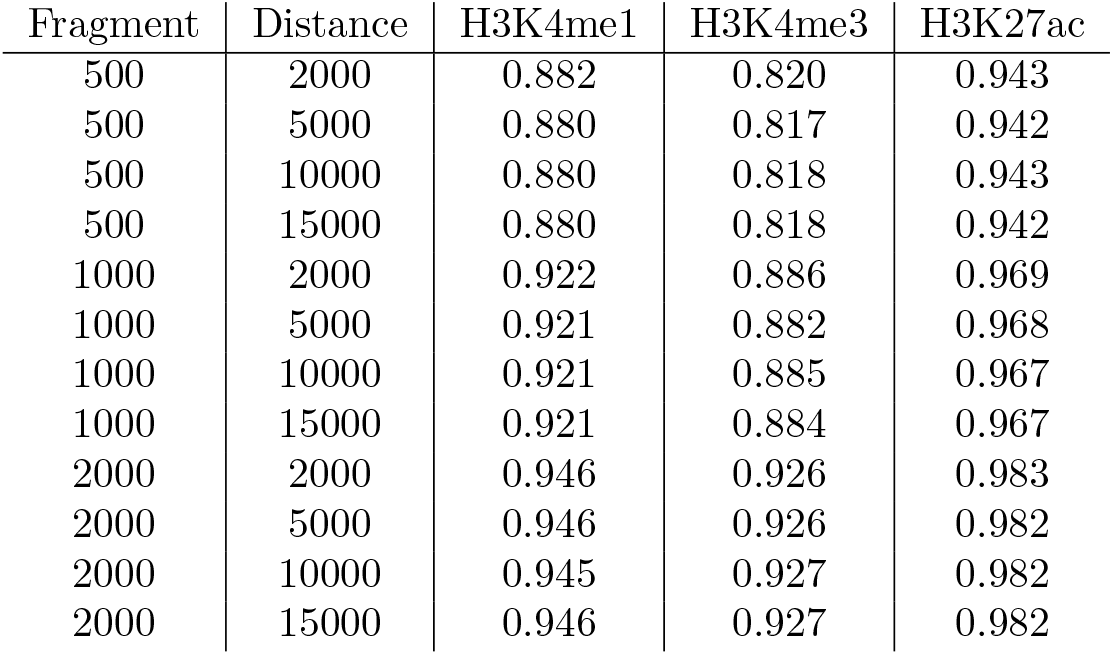
Initial test Regression models (depth 25) First column indicates the fragment size; Second column the distances adjacent to the central fragment; Next columns indicate the Pearson coefficient correlation for the analyzed RFs.

### 2.4 Identification of relationships between PTMs

To determine the relationships between different PTMs, RF regressor algorithms were trained to predict each of the marks by employing different combinations of other marks selected according to the relevance of these marks to predict chromatin structure (Fig 1.A). The idea is to employ the predictability of a mark as a function of combinations of other marks to determine the relationships between them. This is expected to determine four types of relationships: dependency, if a mark is necessary to predict another mark; interdependence - if two marks or combinations of them can better predict another mark, there is an interdependence between the marks used to predict and the target; redundancy - if both one mark and another can predict a third, there is redundancy between them; finally, non-informative relationships, if one mark, or a combination of brands, cannot predict another or worsens its prediction significantly (Fig. 2). A dataset for which both the “target” and input marks are known was used to validate the predictions, so it would be possible to determine when the ML model has been wrong or successful. The results were evaluated using Pearson’s correlation coefficient, where values close to 1 represent a more robust model.

**Figure 2:**
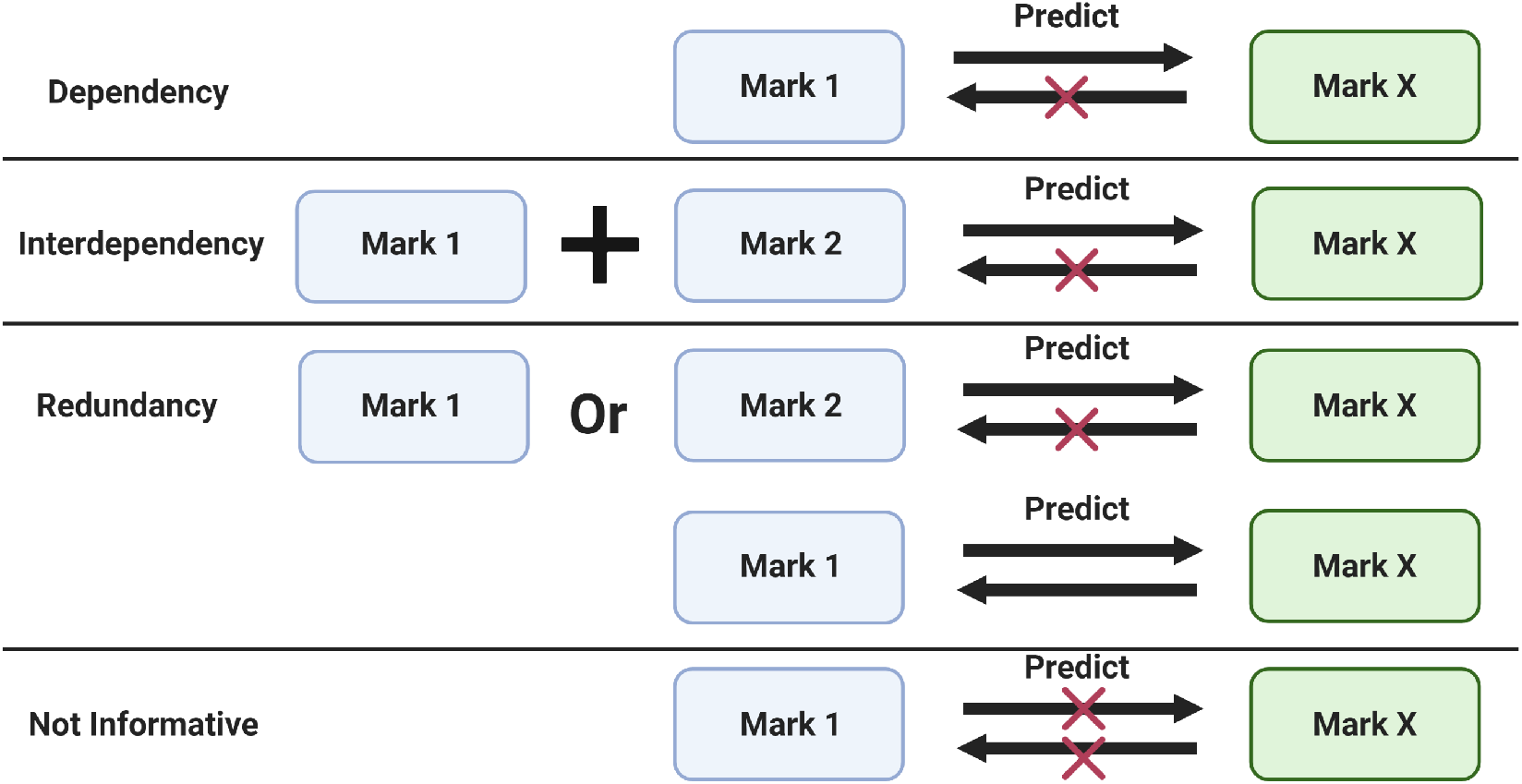
Possible relationships. Dependency, if a mark is necessary for predict another mark; interdependency, if two marks or a combinations of marks are able to predict better another mark; Redundancy, if both mark and another can predict a third there is redundancy between them; not-informative relationships, if a mark, or a combination of marks, cannot predict another

### 2.5 Assignment of chromatin states

We used predicted PTMs as input to train a multi-label RF classifier to assign chromatin states (Fig 1.B). In the first instance, ChromHMM [15] and the experimental data (BAM files) of the 33 PTMs of melanocytes in the two biological conditions (Hmel-noTum and Hmel-Tum) were used to make an assignment of 8 chromatin states. The number of states was based on the work presented in [35]. This assignment was done using fragments of 2000 bases and default parameters. Then, this outcome was used to create a multi-label for the dataset used to train the predictor, where “1” was assigned for state presence and “0” for its absence. In the case of the features, these correspond to predicted PTMs in the fragment. Then, metrics such as Precision (P), which corresponds to the total percentage of elements correctly classified, and Recall (R) the number of elements correctly identified as positives out of the total number of true positives, were used to evaluate the predictor’s performance (See formula below). The name states were labeled, analyzing the enrichment of the PTMs in each state and the presence of genome annotations such as coordinates of promoters, genes, and enhancers. The models were trained using data from chromosome 1 because it is the largest chromosome and it has a greater number of examples. This dataset was divided into five parts of similar size, and five models were trained, using four parts as training and the remaining one as testing. This strategy avoids overfitting by using the exact data for training and testing. Finally, a predictor is obtained to predict each part of the chromosome. In order to predict states in the entire genome, the previously mentioned trained models using datasets for each chromosome were used.

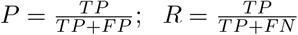

Where a TP (true positive) is a chromatin fragment which state is correctly predicted, a FP (false positive) is a fragment predicted to be in a state different to the one it actually is in, and a FN (false negative) is a fragment that should have been predicted to be in a certain state but it was not.

### 2.6 GRN analysis

The analysis of GRNs was based on the methodology presented in our previous work as explained in Murgas et al. [36], where a reference network is filtered using epigenetic data to remove unlikely regulatory interactions. Here, instead we removed interactions that arise from genes that are in chromatin states known to contain inactive regulatory regions. For this, a human reference network was built using data of promoter coordinates available on the GeneCards database. This database contains information on TFBSs associated with them and the genes they would be regulating. In a first instance, these promoters were filtered based on their distance from the TSSs of a gene, keeping only those that were no more than 5 kb. These promoters were then filtered using the “Active Promoter” chromatin state assigned with the methodology mentioned above. This filter was made considering that the state completely covers the promoter or there are regions associated with this state within the promoter region. The next GRN was carried out with the information of the promoters that would be transcriptionally active, using the TFBSs and the genes that the promoter would be regulating. Therefore, obtaining a Transcription Factor (TF) → target gene network. A final filter was applied, leaving only those TFs whose promoters are in an “Active Promoter” state. In this way, it would be assumed that these TFs would express themselves and exercise their regulation. Finally, to obtain information on the biological processes of selected sets of genes we employed Enrichr [37].

### 2.7 Code availability

All codes employed in this work are freely available under GNU v3 licence at https://github.com/networkbiolab/RF_histonemarks.

## 3 RESULTS

### 3.1 Performance of PTM prediction

First we generated models which use 35 different types of attributes (32 PTMs, and coordinates of Enhancer, Promoter and Gene annotations). These were trained using data from the non-tumorigenic Pmel cell line (Pmel-noTum) followed by a blind test that was performed using data that was not considered in the training stage of the models made of data from non-tumorigenic Hmel cell line (Hmel-noTum), and the tumorigenic Pmel and Hmel cell lines (Pmel-Tum and Hmel-Tum)), so we can observe how the models manage to generalize on this specific problem. With these analyses, it was possible to obtain robust models for the 33 PTMs (positive correlation values above 0.5), where in the case of testing with the second non-tumorigenic cell line (Hmel-noTum), the lowest correlation value was 0.75 for H3K27me3 and the highest for H2BK5ac with 0.97. In the case of the Hmel-Tum cell line, the lowest value was 0.73 for H4K20me2 and the highest for H3K79me1 with 0.96, for Pmel-Tum, H3K4me2 obtained the lowest with 0.69 and the highest H2BK5ac with 0.93 (see Table 3 (35 attr), for the other PTMs, see Supplementary Table 2).

**Table 3:**
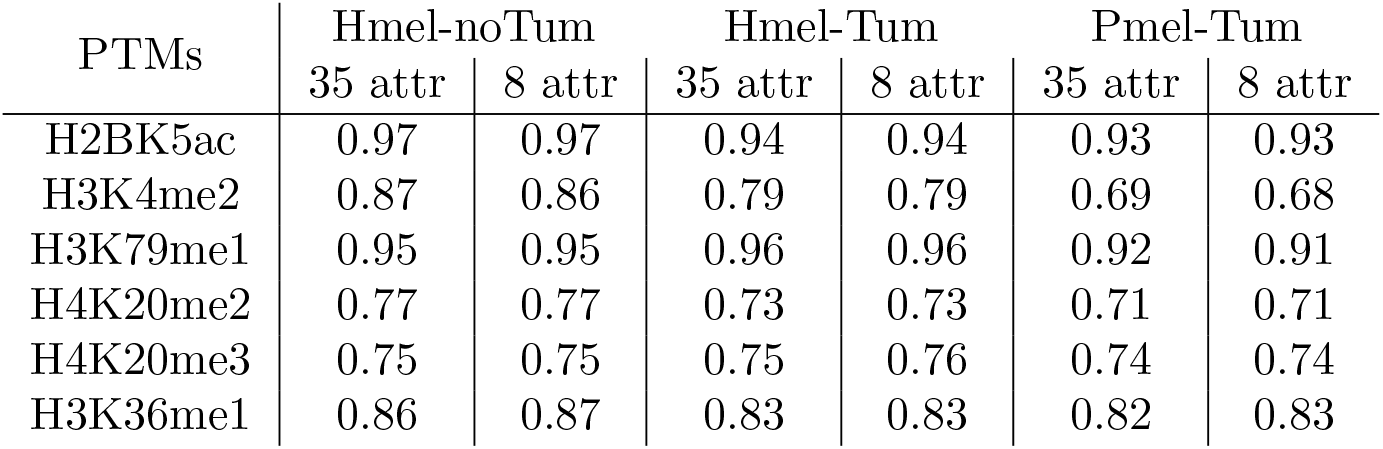
Summary of the results obtained when analyzing the 33 available histone marks with 35 and 8 attributes. These models were trained using the non-tumorigenic cell line Pmel with 35 attributes (32 histone PTMs and annotation data) and 8 attributes with greater importance according to the RFs trained for each mark. The models were tested using the data from the non-tumorigenic cell lines Hmel (Hmel-noTum) and the tumorigenic cell lines Hmel and Pmel (Hmel-Tum and Pmel-Tum). The performance of the models was analyzed using Pearson’s correlation coefficient.

| PTMs | Hmel-noTum |  | Hmel-Tum |  | Pmel-Tum |  |
| --- | --- | --- | --- | --- | --- | --- |
|  | 35 attr | 8 attr | 35 attr | 8 attr | 35 attr | 8 attr |
| H2BK5ac | 0.97 | 0.97 | 0.94 | 0.94 | 0.93 | 0.93 |
| H3K4me2 | 0.87 | 0.86 | 0.79 | 0.79 | 0.69 | 0.68 |
| H3K79me1 | 0.95 | 0.95 | 0.96 | 0.96 | 0.92 | 0.91 |
| H4K20me2 | 0.77 | 0.77 | 0.73 | 0.73 | 0.71 | 0.71 |
| H4K20me3 | 0.75 | 0.75 | 0.75 | 0.76 | 0.74 | 0.74 |
| H3K36me1 | 0.86 | 0.87 | 0.83 | 0.83 | 0.82 | 0.83 |

In the following analysis, we generated models for the different PTMs with a lower number of attributes to test if it is possible to generate robust models with a smaller amount of information. One of the advantages of the RF algorithm is that it allows us to identify which attributes are the most relevant for the prediction. We chose to train new models with the top eight attributes with the greatest importance (as reported by the RF) trained for each PTM. The relevance of all attributes is reported in Supplementary Table 3. The new models showed correlation values similar to those obtained with all the information (32 histone marks, Enhancers, Promoters, and Gene coordinates). For example, in the Hmel-noTum, for H2BK5ac, the same correlation coefficient of 0.97 was maintained with all and only the top 8 attributes; in the case of H3K4me2, it decreased from 0.87 to 0.86, and for H3K36me1 it increased from 0.86 to 0.87 (see Table 3, for a complete comparison for all histone PTMs see Supplementary Table 2).

Next, we inspected our predictions and compared them with actual data. To do so, the BEDgraphs files of the experimental data and the predictions obtained with the 35- and 8-attribute models using chromosome 1 were loaded into the IGV genome browser [38]. Figure 3 shows no significant variations when comparing the predicted data against the real data since the peaks are similar in the entire chromosome. Also, the enrichment peaks were observed at a higher resolution by analyzing the PSMB2 gene, which corresponds to a housekeeping gene found on chromosome 1. As is known, H3K4me3 is mainly enriched in promoter regions, which can be observed when analyzing in the genome browser both for real data and for those predicted in both models (Figure 3.B).

**Figure 3:**
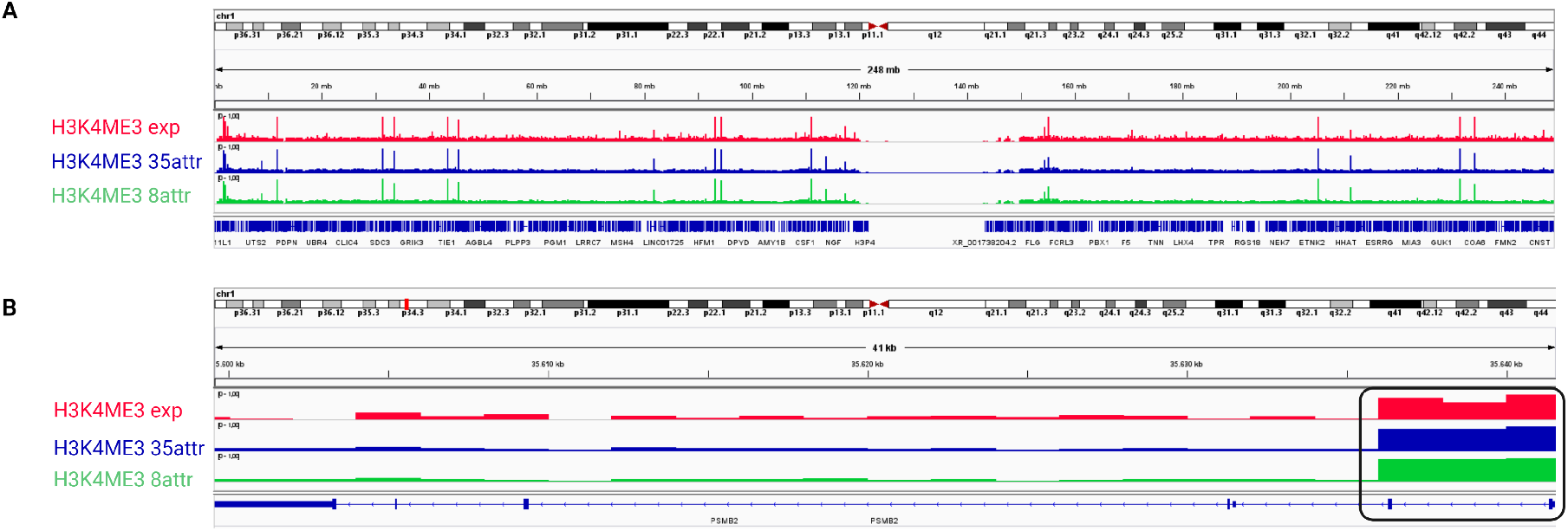
Comparison of peaks of H3K4me3 with IGV genome browser. (**A**) Entire chromosome 1 visualization of the enrichment peaks for H3K4me3, in red the peaks obtained with experimental data, in blue those predicted with the model that uses 35 attributes (32 histone marks, enhancer, gene and promoter annotations) and in green those predicted with the model generated with the 8 most important attributes. (**B**) Visualization of the PSMB2 gene, the square indicates regions of greater enrichment of H3K4me3, corresponding to the promoter region of PSMB2.

As the previous tests were performed using information only from chromosome 1, the generated models were tested using information from the entire genome to see if these models could generalize. These results show that the models do not have significant variations on their performance compared to using only one chromosome, maintaining the same performance for some PTMs or displaying small variations of around 1% (see Supplementary Table 4).

### 3.2 PTMs relationships

Following, we focused on the identification of possible relationships between the analyzed histone PTMs, using the 8 most important attributes reported by the RF models. We performed a manual grouping where those PTMs that shared three attributes in their respective top 8 were grouped together. For group 11, three shared attributes could not be detected, only two. In this way, 12 groups were identified (see Table 4).

**Table 4:** Groups of PTMs that share attributes. First column indicates the groups made based on the shared marks that predicts them better second column the PTMs belonging to each group and the third column the shared attributes for each group. The groups share three PTMs except for group 11 which shares only two.

| Groups | PTMs | Shared attributes |
| --- | --- | --- |
| 1 | H3K23ac, H3K4ac | H3K27ac, H3K14ac, H4K5ac |
| 2 | H3K9me1, H3K36me3, H4K8ac, H3K27me1 | H4K12ac, H3K4ac, H4K16ac |
| 3 | H3K4me1, H2AK5ac, H4K91ac, H4K5ac, H2BK5ac, H3K36ac | H3K4ac, H3K27ac, H2BK120ac |
| 4 | H3K79me2, H3K27me3, H4K20me1 | H3K79me1, H3K79me3, H4K12ac |
| 5 | H3K36me2, H3K36me1 | H3K14ac, H3K27me3, H4K20me1 |
| 6 | H3K27ac, H3K9ac, H3K14ac | H3K4ac, H3K23ac, H3K4me2 |
| 7 | H2BK120ac, H3K18ac | H2BK5ac, H3K4ac, H3K4me1 |
| 8 | H3K4me2, H3K4me3 | H3K9me3, H3K4me1, H3K9ac |
| 9 | H4K20me3, H4K20me2, H3K79me3 | H3K4ac, H3K9me3, H4K20me1 |
| 10 | H3K79me1, H3K9me3 | H4K16ac, H4K20me1, H3K36me2 |
| 11 | H2BK15ac, H4TETRAac | H3K4ac, H3K18ac |
| 12 | H4K16ac, H4K12ac | H4K8ac, H3K79me1, H4K5ac |

This grouping of histone PTMs indicated that it would be possible to predict any mark of those belonging to each group using the same attributes. For example, using the shared attributes of group 1 (H3K27ac, H3K14ac, and H4K5ac), it would be possible to predict any of the PTMs belonging to group 1 (H3K23ac and H3K4ac). To verify this, new models were generated in which only these shared attributes were used to predict each group different PTMs. These models show robust performance values for all predicted PTMs. For example, when analyzing group 3, which is the one with the highest number of PTMs, Pearson’s correlation values between 0.87-0.96 are obtained when tested in the Hmel-noTum, values between 0.82-0.94 when tested in the Hmel-Tum and values between 0.78-0.93 when tested in the Pmel-Tum (Table 5, for the other groups see Supplementary Table 5).

**Table 5:** Models obtained for group 3, using only three shared attributes to predict each mark. The models were trained using the non-tumorigenic cell line Pmel (Pmel-noTum) with the three shared attributes of group 3, the models were tested using the data from the non-tumorigenic cell lines Hmel (Hmel-noTum) and the tumorigenic cell lines Hmel and Pmel (Hmel-Tum and Pmel-Tum). The performance of the models was analyzed using Pearson’s correlation coefficient

| PTMs | Hmel-noTum | Hmel-Tum | Pmel-Tum |
| --- | --- | --- | --- |
| H3K4me1 | 0.87 | 0.82 | 0.83 |
| H2AK5ac | 0.91 | 0.88 | 0.88 |
| H4K91ac | 0.94 | 0.90 | 0.78 |
| H4K5ac | 0.94 | 0.91 | 0.87 |
| H2BK5ac | 0.96 | 0.94 | 0.93 |
| H3K36ac | 0.91 | 0.85 | 0.82 |

Additionally, new models were trained to determine relationships between PTMs, varying the number of attributes, using only the most important attribute, and increasing the number of attributes until reaching the eight most important ones for each mark.

In the models for the 33 PTMs, it is observed that, in general, all except for H3K27me3 and H3K4me3 present positive correlation values above 0.50 in all the cell lines evaluated using only the most important attribute. There is an improvement in performance for the prediction of all histone marks by adding the second most important attribute, which would indicate possible interdependence relationships. Adding the third attribute we observed a drastic decrease for most of the PTMs evaluated, except for the models for H2BK120ac, H3K4ac, and H4K91ac, for which the performance improvement or is maintained at the same level. In the case of adding the fourth attribute, improvements are observed compared to the previous models, but in most cases, the correlation coefficients obtained are not greater than those obtained with the models of the two most important attributes. Finally, improvements are observed in some of the cell lines evaluated from the fifth attribute, which begin to have slight variations when adding more attributes. A detailed table with results can be seen in Supplementary Table 6.

### 3.3 Comparison with other methods

We next compared our approach against two methodologies for predicting PTMs from epigenetic data. These methods correspond to CHROMIMPUTE, which uses a methodology based on regression-type decision trees, and AVOCADO, which uses a tensor factorization approach. For both comparisons, Pmel-noTum and Hmel-noTum on chromosome 1 were used, as in our method’s training and testing stage. For each compared model, the same histone marks used in our method were used as attributes, predicting the mark in the Hmel-noTum cell line.

In the case of CHROMIMPUTE, it accepts the same BEDgraphs used to test our method, so the comparison is straightforward. For AVOCADO, the data needed to be reanalyzed with the same processing pipeline as our method but using 250-base fragments due to the input format required by this tool. Since our method uses 2000-base fragments we needed to transform the output of AVOCADO as follows: the average predicted signal was calculated for every eight fragments and used as the label for the equivalent 2kb BEDgraphs.

We first compared the models that use all the available histone PTMs information. The performance of each method was evaluated by calculating Pearson’s correlation between the predicted data against the experimental data. Additionally, to corroborate that the differences between the performance all the compared methods were significant or not, 80% of the predicted and experimental data were randomly selected, and a new Pearson correlation was calculated; This process was performed 100 times for each PTM to apply a t-test to determine whether the differences are significant or not (Supplementary Table 7). Using all PTMs, our approach shows a better performance than CHROMIMPUTE for the 33 PTMs analyzed, reaching a very large difference in PTMs such as H3K27me3 where our approach has a correlation coefficient of 0.81 and CHROMIMPUTE of only 0.32. In the case of the comparison with AVOCADO, the latter had better performance in three of the 33 PTMs analyzed (H3K27me1, H3K27me3, and H4K20me2). Regarding the 8-attribute models, the same scenario is repeated when comparing our method against CHROMIMPUTE, showing worse performance for all marks with with the latter. When comparing against AVOCADO, this other approach is only better that the 8-attributes models for PTM (H3K27me3). Finally, CHROMIMPUTE also fails to outperform our approach when using our models with fewer attributes. In the case of AVOCADO, it showed better performance than our models with less than eight attributes for four PTMs (H3K23ac, H3K27me3, H3K79me1, and H3K79me3). These results show that our methodology is better than CHROMIMPUTE regardless of the number of attributes used to generate the model. In the case of AVOCADO, this method is only better than our approach in few particular cases. It should be noted that the performances obtained by our method when using all PTMs are very similar to those generated using only the 8 most important attributes and that these are surpassed only in three PTMs predicted with AVOCADO (H3K27me1, H3K27me3, and H4K20me2) (see Table 6 and Supplementary Table 8 for a detailed comparison mark by mark).

**Table 6:** Comparison of our Random Forest approach (RF) with CHROMIMPUTE and AVOCADO. The first column indicate the PTM predicted. The next columns show the results of the generated models using different amount of attributes. The performance of the models was analyzed using Pearson’s correlation coefficient. In bold number indicate the method with statistically significant better results using a t-test (p-value *≤* 0.05).

| PTM | AllAttr |  |  | 8Attr |  |  | 3Attr |  |  |
| --- | --- | --- | --- | --- | --- | --- | --- | --- | --- |
|  | RF | CHROMIMPUTE | AVOCADO | RF | CHROMIMPUTE | AVOCADO | RF | CHROMIMPUTE | AVOCADO |
| H3K27me3 | 0.81 | 0.32 | **0.83** | 0.82 | 0.35 | **0.83** | 0.55 | 0.3 | **0.8** |
| H3K27me1 | 0.83 | 0.76 | **0.88** | **0.83** | 0.74 | 0.79 | **0.84** | 0.78 | 0.76 |
| H4K20me1 | **0.85** | 0.73 | 0.64 | **0.85** | 0.72 | 0.63 | **0.77** | 0.63 | 0.74 |
| H4K20me2 | 0.77 | 0.63 | **0.79** | **0.77** | 0.66 | 0.7 | 0.72 | 0.57 | 0.72 |
| H3K23ac | **0.92** | 0.84 | 0.88 | **0.92** | 0.85 | 0.87 | 0.86 | 0.78 | **0.87** |
| H3K79me1 | **0.95** | 0.9 | 0.81 | **0.95** | 0.88 | 0.86 | 0.74 | 0.6 | **0.8** |
| H3K79me2 | **0.95** | 0.86 | 0.8 | **0.95** | 0.88 | 0.89 | **0.95** | 0.88 | 0.92 |
| H3K79me3 | **0.88** | 0.83 | 0.86 | **0.88** | 0.83 | 0.86 | 0.72 | 0.6 | **0.79** |
| H3K4me3 | **0.88** | 0.81 | 0.74 | **0.88** | 0.8 | 0.75 | **0.75** | 0.69 | 0.5 |
| H3K9me3 | **0.78** | 0.48 | 0.73 | **0.78** | 0.52 | 0.69 | 0.69 | 0.29 | 0.69 |
| H3K18ac | **0.96** | 0.89 | 0.93 | **0.95** | 0.89 | 0.91 | **0.94** | 0.91 | 0.91 |

### 3.4 Test in Colorectal Cancer

We next tested our approach on ChIP-seq data generated from colon carcinoma cells (HCT116) for ten PTMs (H3K27ac, H3K27me3, H3K36me3, H3K4me1, H3K4me2, H3K4me3, H3K79me2, H3K9ac,H3K9me3 and H4K20me1). This data was obtained from the ENCODE database and we re-analyzed it with to the pipeline used in this work. Based on the feature performance obtained with the previously explained models and the models trained with three to eight marks, we predicted the remaining 23 PTMs using a cascade of predictors (Figure 4).

**Figure 4:**
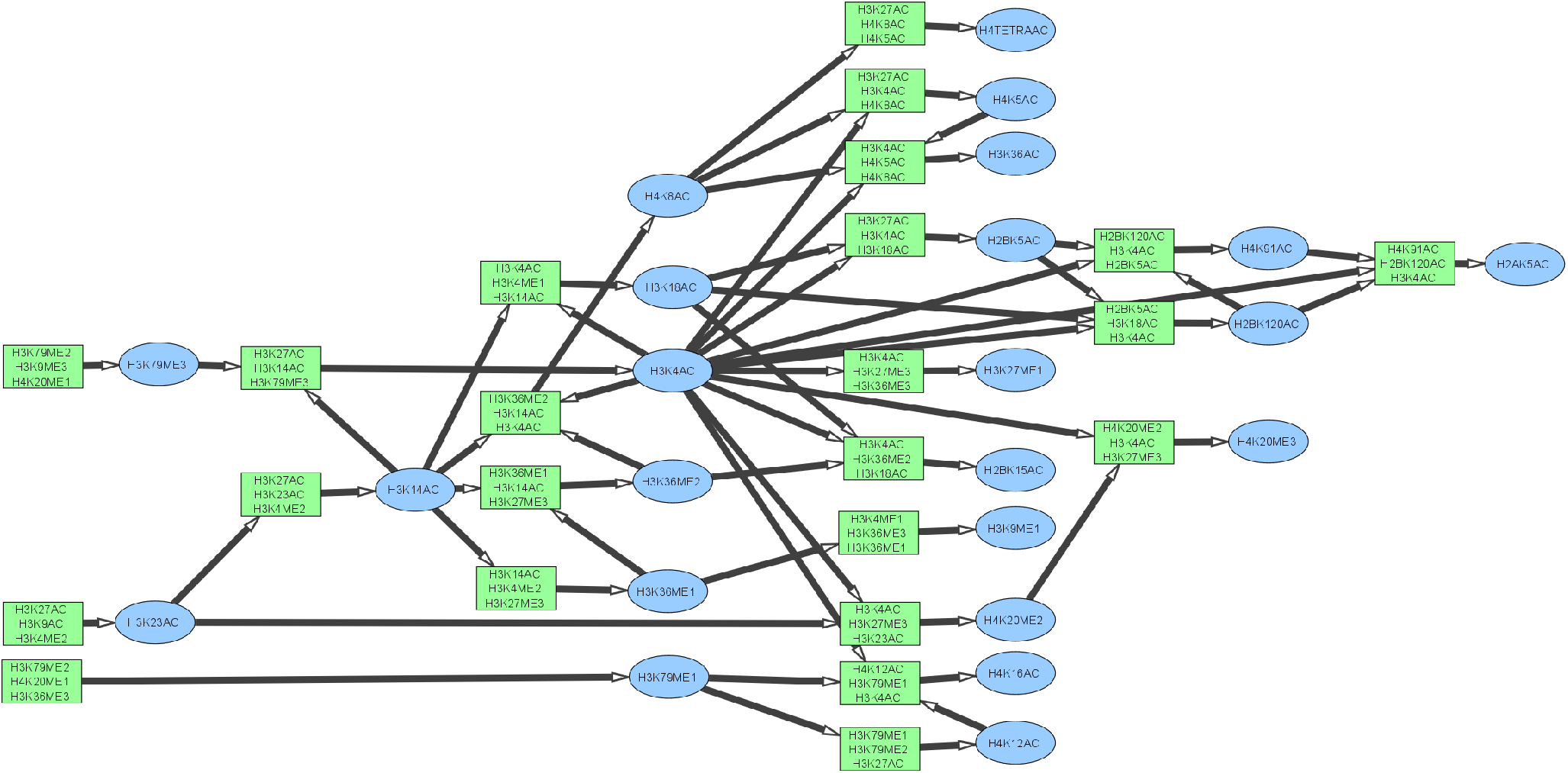
Theoretical analysis of the prediction of the 22 histone PTMs. Using the colon dataset that contains 11 PTMs (H3K27ac, H3K27me3, H3K36me3, H3K4me1, H3K4me2, H3K4me3, H3K79me2, H3K9ac, H3K9me2, H3K9me3 and H4K20me1) and using three of the eight most important attributes (squares), the different PTMs analyzed in this work (ovals) are predicted.

In the first instance, we generated models to predict four of the histone marks from the colon cancer dataset (H3K4me1, H3K4me2, H3K4me3 and H3K9ac). To train these models we used data from the non-tumorigenic Pmel cell line for chromosome 1 and tested it with the non-tumorigenic Hmel and the two tumorigenic cell lines on the same chromosome. Additionally, the model was also tested using whole genome information for the colon carcinoma cell line (HCT116). The results show robust models for the four marks analyzed in all the cell lines analyzed (Table 7), except for H3K4me3, which in Pmel-noTum show values below 0.5. It should be noted that when testing in the HCT116 cell line, it yields correlation values above 0.9, indicating that the models generalize in different cell lines, including different tumor lines where the epigenome may follow different patterns (Table 7).

**Table 7:** Models trained with non-tumorigenic Pmel and tested on different cell lines including colon carcinoma. The models were trained using the non-tumorigenic cell line Pmel with the three histone marks present in carcinoma dataset, the models were tested using the data from the non-tumorigenic cell lines Hmel (Hmel-noTum), the tumorigenic cell lines Hmel and Pmel (Hmel-Tum and Pmel-Tum) and HCT116 cell line. The performance of the models was analyzed using Pearson’s correlation coefficient.

| PTMs | Hmel-noTum | Hmel-Tum | Pmel-Tum | HCT116 |
| --- | --- | --- | --- | --- |
| H3K4me1 | 0.86 | 0.80 | 0.79 | 0.92 |
| H3K4me2 | 0.86 | 0.77 | 0.66 | 0.97 |
| H3K4me3 | 0.86 | 0.75 | 0.45 | 0.95 |
| H3K9ac | 0.87 | 0.79 | 0.85 | 0.92 |

Additionally, an analysis similar to the previous one was performed but this time using data from HCT116 the cells to train the models. These models were trained using the chromosome 1 data from HCT116 and tested on the same chromosome on the other cell lines but on HCT116, were we employed chromosomes 2 to 22. The results obtained show a slight decrease when testing the model in cell lines that are not colon carcinoma, but an increase in performance is seen when testing with the latter (Table 8).

**Table 8:** Models trained with HCT116 and tested on different cell lines including colon carcinoma. The models were trained using the HCT116 cell line with the three PTMs present in carcinoma dataset, the models were tested using the data from the non-tumorigenic cell lines Hmel (Hmel-noTum), the tumorigenic cell lines Hmel and Pmel (Hmel-Tum and Pmel-Tum) and HCT116 cell line. The performance of the models was analyzed using Pearson’s correlation coefficient.

| PTMs | Hmel-noTum | Hmel-Tum | Pmel-Tum | HCT116 |
| --- | --- | --- | --- | --- |
| H3K4me1 | 0.81 | 0.72 | 0.65 | 0.97 |
| H3K4me2 | 0.84 | 0.83 | 0.65 | 0.98 |
| H3K4me3 | 0.86 | 0.73 | 0.46 | 0.97 |
| H3K9ac | 0.83 | 0.80 | 0.83 | 0.93 |

### 3.5 Prediction cascade

To see if it was possible to predict histone PTMs using predicted data instead of ChIP-seq experimental data, and thus, to reduce the number of experiments necessary for predicting each mark, we used ten experimentally determined histone PTMs of Pmel-noTum based on the list of PTMs available for colon cancer (HCT116) and trained models following the cascade of predictors mentioned above (Fig. 4). The performance of predictors was tested using a dataset with experimental data of Hmel-noTum and models were trained only on predicted data and experimental data from those ten initial marks following a rigorous separation between training and test data to avoid overestimating performances. The results show that the number of histone marks available for a sample can be increased from only a subset of ten experimentally determined marks. We see a slight decrease in the performance of predictions using only experimental data compared to the mixed data. For example, with H2BK5ac, we observe a decrease from 0.97 to 0.95. On the other hand, for both H4K8ac and H4K12ac, the performance obtained was the same in both cases. Finally, the mark with the worst performance was H3K36me2, with a value of 0.64.

### 3.6 Chromatin state assignment using predictions

The following analysis consisted of using data from PTMs predicted with the proposed methodology for assigning chromatin states. A gold standard was first generated using the ChIP-seq data from the 33 histone PTMs of Hmel-noTum and the ChromHMM tool to assign 8 chromatin states (“poised enhancer”, “active enhancer”, “weak promoter”, “active promoter”, “poised promoter”, “transcriptional elongation”, “repressive state” and “low signal state”) which were assigned according to the enrichment of PTMs in each state, based on [35]. Because the generated RF models predict peaks in BEDgraph format, it is impossible to use these data directly with the ChromHMM software. For this, a new predictive model based on multi-label RF classifier was generated. Two models were trained using the predicted data for all 33 histone marks and for only ten marks (H3K27ac, H3K27me3, H3K36me3, H3K4me1/me2/me3, H3K79me2, H3K9ac, H3K9me3, and H4K20me1). As target label we employed the chromatin states generated with ChromHMM using the experimental data of the 33 PTMs. This subset of ten marks was selected because they are among the most characterized in databases, and some are of clinical interest due to their association with cancer [39, 40]. These models can generate robust predictions of chromatin states using only predicted data. Precision (P) and Recall (R) values ≳ 70% are obtained for both models on the genome not used to train the predictors (chromosomes 2 to 22 and X), with the worst performance was for the state “Poised enhancer” state (see Table 10).

**Table 9:** Predictor’s cascade. Performance of Predictor cascade. 23 histone marks were predicted using a cascade Predictors from experimental data from ten histone PTMs. The fist column indicate the predicted PTMs; the second column show the performance using experimental only experimental data to test; the third column show the performance obtained with mixed data (using experimental and predicted data or in some case only predicted data). The performance of the models was measured using Pearson’s correlation against actual mark data.

| PTM | Hmel-noTum exp | Hmel-noTum mixed |
| --- | --- | --- |
| H2AK5ac | 0.92 | 0.83 |
| H2BK120ac | 0.94 | 0.89 |
| H2BK15ac | 0.74 | 0.71 |
| H2BK5ac | 0.97 | 0.95 |
| H3K14ac | 0.87 | 0.82 |
| H3K18ac | 0.93 | 0.91 |
| H3K23ac | 0.85 | - |
| H3K27me1 | 0.80 | 0.77 |
| H3K36ac | 0.90 | 0.88 |
| H3K36me1 | 0.80 | 0.72 |
| H3K36me2 | 0.83 | 0.64 |
| H3K4ac | 0.92 | 0.90 |
| H3K79me1 | 0.95 | - |
| H3K79me3 | 0.86 | - |
| H3K9me1 | 0.88 | 0.86 |
| H4K12ac | 0.90 | 0.90 |
| H4K16ac | 0.81 | 0.78 |
| H4K20me2 | 0.73 | 0.68 |
| H4K20me3 | 0.74 | 0.65 |
| H4K5ac | 0.95 | 0.92 |
| H4K8ac | 0.85 | 0.85 |
| H4K91ac | 0.94 | 0.86 |
| H4TETRAac | 0.92 | 0.89 |

**Table 10:**
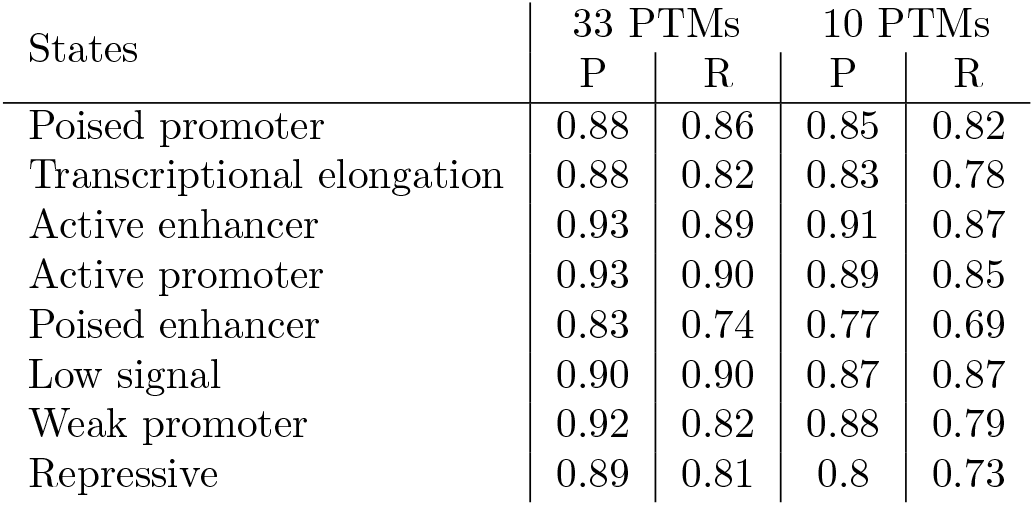
Chromatin states assignment using predicted PTMs. 8 chromatin states were assigned by means of a classifier RF-based predictor, which uses predictions obtained by our random forest models and labels obtained with ChromHMM as input data. State labels were assigned according to the enrichment of 33 histone marks experimentally determined. The performance of the models was measured using to the Precision (P) and Recall (R) metrics.

| States | 33 PTMs |  | 10 PTMs |  |
| --- | --- | --- | --- | --- |
|  | P | R | P | R |
| Poised promoter | 0.88 | 0.86 | 0.85 | 0.82 |
| Transcriptional elongation | 0.88 | 0.82 | 0.83 | 0.78 |
| Active enhancer | 0.93 | 0.89 | 0.91 | 0.87 |
| Active promoter | 0.93 | 0.90 | 0.89 | 0.85 |
| Poised enhancer | 0.83 | 0.74 | 0.77 | 0.69 |
| Low signal | 0.90 | 0.90 | 0.87 | 0.87 |
| Weak promoter | 0.92 | 0.82 | 0.88 | 0.79 |
| Repressive | 0.89 | 0.81 | 0.8 | 0.73 |

Additionally, we generated a new model to predict chromatin states, but this time using the 23 PTMs predicted marks of Hmel-noTum obtained with the cascade of predictors and the experimental data of the remaining ten PTMs. This new model shows P and R values ≳ 70% in seven states. Only the state “Poised enhancer” obtained an R value of 57%. To test if the use of predictions plus experimental data improves the performance of chromatin assignment, a new model was generated that use only the experimental data from the ten PTMs used for the previous model. When comparing the performance of both models, it was observed that including predicted data improves the chromatin assignment in all states (Table 11).

**Table 11:** Chromatin states assigned with predictors cascade. Chromatin states assignment using cascade predictors and experimental data. 8 chromatin states were assigned by means of a classifier RF-based predictor, which uses predictions obtained by our random forest models cascade and labels obtained with ChromHMM as input data. A second assignment was made using only experimental data of ten histone marks. State labels were assigned according to the enrichment of of 33 histone marks experimentally determined. The performance of the models was measured using to the Precision (P) and Recall (R) metrics.

| State | PTMs cascade |  | 10 exp PTMs |  |
| --- | --- | --- | --- | --- |
|  | P | R | P | R |
| Poised promoter | 0.84 | 0.81 | 0.81 | 0.74 |
| Transcriptional elongation | 0.78 | 0.7 | 0.77 | 0.68 |
| Active enhancer | 0.89 | 0.81 | 0.89 | 0.79 |
| Active promoter | 0.85 | 0.78 | 0.83 | 0.75 |
| Poised enhancer | 0.71 | 0.57 | 0.69 | 0.49 |
| Low signal | 0.87 | 0.9 | 0.84 | 0.89 |
| Weak promoter | 0.9 | 0.8 | 0.85 | 0.73 |
| Repressive | 0.91 | 0.86 | 0.9 | 0.82 |

### 3.7 Case study: Colorectal cancer

To test the utility of our methodology, we decided to use colorectal cancer cell line HCT116 as a case study model. In the first instance, we use the ChIP-seq data of the ten PTMs from HCT116 (H3K27ac, H3K27me3, H3K36me3, H3K4me1, H3K4me2, H3K4me3, H3K79me2, H3K9ac, H3K9me3, and H4K20me1) to predict the 23 remaining PTMs and increase the number of marks available for these samples. These predictions were made using the cascade of predictors implemented in this work. The next step was to use this data to predict the chromatin states assignment using the previously trained predictor that uses the predictions obtained with the cascade of predictors. In this way, we obtained an assignment of chromatin states of cell line HCT116 for the whole genome. To observe the biological importance of the results and validate the use of predicted PTMs we carried out an analysis of GRNs to study how the regulation of specific genes is linked to chromatin states. Here we constructed GRNs for the predicted colorectal chromatin states. The methodology used was based on Murgas et al. [36], which filters a reference network from epigenetic information, leaving only those connections occurring in contexts where the regulation can take place according to what we know on the relationship between epigenetics and TF activity. In our case, the reference network was built using the promoter information obtained in the GeneCards database [29], using data of TFBSs and gene targets of promoters, and filtering it by those states that would be transcriptionally active. According to the assignment of states made with the predictions of PTMs, specifically in this case, using the state of “Active Promoter”. Given the large size of the complete GRN, we focused here on the subnetwork that explains changes for genes in the chromosome 1. The resulting GRNs has 7342 edges, where 64 nodes correspond to TFs and 252 to target genes. We next characterized the functions of target genes using the tool Enrichr [37] to determine enriched “biological process” GO terms. With these analyses we found enrichment in processes associated to DNA damage such as “mitotic G1 DNA damage checkpoint signaling”, “DNA damage response”, among others (Supplementary Table 9). Additionally, analyzing the list of target genes obtained in the GRN of chromosome 1, eight genes were found previously associated to colorectal cancer (IL6R, KISS1, LEPR, LMNA, MUC1, PER3, PTGS2, and TACSTD2). The same analysis but for the whole genome network is described in Supplementary Data.

## 4 DISCUSSION

Epigenetic marks are usually characterized to determine the activity of *cis* and *trans* regulatory elements in the genome. For this, several marks have been associated to active or repressive promoter activity [2], or similarly for the activity of enhancers [41] or for the binding of TFs to the chromatin [42, 43]. Another application of these marks is to determine chromatin states, a more detailed definition that subdivides the traditional euchromatin/heterochromatin into several functional states, that is known to require many different marks for consistency [44]. One of the less studied aspects of epigenetic marks is the existence of relationships between different marks where the effect of a single histone PTM is altered by the presence of other marks nearby [45, 46]. Even more important would be to demonstrate the existence of an epigenetic code where different patterns of epigenetic modifications of histones linked to specific regions would be associated to specific effects [7, 8].

In this work, we describe a new tool for the prediction of epigenetic marks from other epigenetic PTMs. Our approach allows us to increase the amount of data available for studying different tissues or cell lines for which there is not much data available, aiming to reduce the cost of analyses that require many histone marks. Moreover, our approach, based on RFs, allowed us to establish several relationships of interdependency and redundancy between different epigenetic marks, helping in this way to provide evidence towards the establishment of the human epigenetic code. We chose RFs as our predictive algorithm because in addition to accurate predictions when the dataset allows it, it reports a ranking of importance for each attribute used as input [34].

When comparing our RF predictor with other with existing methodologies, the most relevant difference is that the RF generates interpretative relationships that for example, neural networks cannot generate. In absolute terms, CHROMIMPUTE is worse for all marks than our tool, while AVOCADO is better for only a few marks independently of the number of marks employed as input. These results indicate that despite the algorithm used, our RF models are out performing other existing methods, and in addition, their modest requirements in terms of hardware make our models more readily usable. The capability to determine the relevance of the input attributes is a very important difference with other algorithms since, in this application at least, it is as important to understand the relationships reported when training the RFs as to obtain good predictions.

Given the output format of our methodology, which consists of BEDgraphs files, it was not possible to directly use tools to assign states to chromatin, such as ChromHMM. This incompatibility is something that can be improved in the future for our method and for the state assignment methodologies because many of the epigenetic data available in public databases are in this format, making it impossible to use them for tasks of this type. That is why, to verify if the use of predictions obtained with our method, the assignment of chromatin states was possible, a state predictor was made using as target the states generated by ChromHMM and as predicted BEDgraphs for histone marks. This model achieved robust states assignment very similar to that obtained with experimental data (Tables 10).

We have applied our RF approach to reduce the number of histone modifications required to robustly annotate chromatin states.

Generally, it is thought that greater precision is expected by having grater amount of data because more information about the problem to be solved would be available. However, our results show that it is possible to generate predictions of equal or better quality using a smaller number of experiments (as shown in Tables 3, 5). Importantly, this would also support the existence of dependency and redundancy relationships among the PTMs. For example, when avoiding the use of a certain mark but using other marks to make predictions, if the results are not altered we can assume that there is redundancy between the marks used and the one not used to train the predictor. When the use of a mark worsens the performance of the prediction of another mark we can assume there is no relationship between them or that it is counter informative. Similarly, if more than one mark is required to predict accurately another mark and without these marks, or using them separately, the predictions are much worse, we assume there is an interdependence relationship. This relationship is observed, for example, with the analysis of H4K16ac, a mark related to breast, prostate, and colorectal cancer [39, 40], the results of which show a relationship of the interdependence of H4K12ac and H4K8ac (Supplementary Table 3 and 6). Importantly, the existence of these relationships support the existence of the epigenetic code [7, 8].

Another type of relationship is defined by grouping histone PTMs using shared attributes from those that are among the most important for their prediction. In this way, redundancy-type relationships were identified. For example, we could say that H3K23ac and H3K4ac, which belong to group 1 (Table 4), would have a redundancy relationship with H3K27ac, H3K14ac, and H4K5ac since they all share the three marks as predicting attributes. Importantly, we generated robust models only using these three marks to predict H3K23ac and H3K4ac (Supplementary Table 5). Thus, given that it is possible to determine where two marks are present by determining the other three, our predictions provide some evidence that the epigenetic code is redundant.

We also observed a slight drop in the performance of the models trained on non-tumoral cells when tested on tumor cells. This phenomenon may be because there is a known alteration in the patterns of certain marks in tumors [22]. Therefore, the cascade of alterations caused at many levels in tumors affects the way epigenetic marks are related to each other and could affect our predictions [47, 48]. Our results also indicate that at least in the cell lines we employed in this study, there are some marks with prediction results that are not different between tumoral and non-tumoral cells, as indicated by the good generalization capabilities of our models (Table 4). These findings suggests that only the levels and location of certain marks are affected and that these changes only affect partially the relationships found using our RF approach.

Next, we analyzed if our models were biased to the cell lines used in their training in which it was being analyzed. For this test we employed a dataset of 10 histone marks form the HCT116 cell line, which corresponds to colon carcinoma. These analyses indicated good results for those models trained with Pmel-noTum data on HCT116 (Table 7). The only exception was observed for H3K4me3 when testing on Pmel-Tum, which could be because the marks with the most significant relationships with it are not in this test data. Importantly, this test on a different cell line validates the generalization capabilities of our approach since it works largely independently of the cell line used.

Our additional analysis of GRNs based on these predicted chromatin states using colorectal cancer cell line as case study, confirmed the validity of our results based on the functions linked to genes target obtained with the filters performed. For example, among these genes we found PTGS2. This gene has been associated with inflammatory stimuli and expressed in tumor cells of colorectal cancer [49, 50]. Another example is VEGF-A, a gene that encodes an angiogenic factor that has been detected up-regulated in different types of tumors such as colorectal cancer [51, 52]. These two genes, already known to be linked to tumors and those others linked to other cancer related functional terms indicate that despite using a reduced amount of experimentally determined histone marks that where used to generate predictions of other marks for the state assignment, the resulting GRNs support the valid of our whole approach.

## 5 CONCLUSION

We have described here a new approach based on RFs to predict histone marks from other histone marks. We first proved that our approach is robust and outperforms other available tools that perform the same task. Moreover, given the intrinsic ability of the RF to determine the relevance of each input feature, our method produced robust predictions relying on only a few other marks. Nonetheless, it is even more important that by using our tool is possible to accurately determine many histone marks, increasing the number of histone PTMs available from the same sample and thus, allowing more analysis from the same experimental data. For example, by combining our approach with ten experimentally determined marks we were able to accurately assign states to chromatin in a way that allowed to determine GRN based on active promoters. This GRN analysis, applied to a colorectal cancer cell line, was validated by the genes whose expression is linked to this type of cancer.

Finally, it is very important to highlight that in the process of creating the cascade of predictors we found evidence to support the existence of a robust and redundant epigenetic code. In this code, we have shown how certain marks are accurately predicted by other marks, i.e., there are redundancies; and some other marks are indispensable to predict the existence of others, which indicate an inter-dependence between marks that in a way also explains the large alterations in the epigenetic profiles that is often observed in complex diseases.

Our approach, simply based on a prediction algorithm would benefit from the availability of more data from other samples or other types of information such as gene expression profiles or DNA accessibility. Nonetheless, our new approach is useful on its own and will surely help other scientists in their work.

## 6 ACKNOWLEDGEMENTS

This work was funded by: ANID PhD fellowship [21201856] to LM, FONDECYT Regular Projects [1181089] to AJMM and [1191526] to ER, FONDECYT Inicio [11171015] to MS, and Centro Ciencia & Vida, FB210008, Financiamiento Basal para Centros Científicos y Tecnológicos de Excelencia de ANID. PoweredNLHPC: this research was partially supported by the supercomputing infrastructure of the NLHPC (ECM-02).

