## Supplementary material for "Random Forest approach for the identification of relationships between epigenetic marks and its application to robust assignment of chromatin states": supplementary_data.pdf

### 1 Supplementary Data

#### 1.1 Post translational modifications (PTMs) of interest in cancer

A more exhaustive analysis was carried out for seven PTMs (H4K16AC, H3K18AC, H3K4ME3, H3K27ME3, H3K36ME2, H4K20ME2/3) that have greater biological relevance since they are known to play an important role in different types of tumors [1] [2]. We trained models by adding marks in growing order of relevance as reported by the Random Forests (RFs) trained with all 35 descriptors.

### 1.1.1 H4K16AC

The model obtained with the most important attribute (H4K12AC) has a correlation value of 0.80 in the Hmel non-tumor condition, 0.79 and 0.84 respectively for Hmel and Pmel tumorigenic. With the addition of the second attribute, a slight increase is observed, obtaining 0.81 in the non-tumor condition and 0.80 in Hmel tumorigenic. Adding the third attribute, a decrease in the correlation coefficient is observed in all conditions, which is increased with the addition of the fourth attribute, but is still lower than the model generated with two attributes. With the fifth attribute, an increase in correlation is seen again, this time obtaining values of 0.81 for the non-tumor condition, 0.81 and 0.85 in Hmel and Pmel tumorigenic, respectively. Finally, adding attributes 6-8 slight variations are observed. When making a model using only the second most important attribute (H4K8AC) to see if there was a dependency relationship or only interdependence, it is observed that it is possible to generate a robust model, but of lower quality than using only the most important attribute. This suggests the presence of a dependency relationship of H4K8AC with H4K16AC, and an interdependence relationship between the first and second para attributes with H4K16AC.

### 1.1.2 H3K18AC

Using only the most important attribute H2BK5AC to predict H3K18AC we obtained Pearson's correlation values of 0.92 for the non-tumor condition and 0.78 and 0.84 respectively for Hmel and Pmel tumorigenic. Adding the second attribute (H2BK120AC) shows an increase in correlation to 0.94 for the non-tumor condition and 0.83 and 0.88 for Hmel and Pmel tumorigenic. This would indicate a possible relationship of interdependence between H2BK5AC and H2BK120AC with H3K18AC. By adding the third most important mark, a decrease is observed in the non-tumor line and Pmel tumorigenic with 0.92 and 0.86, in the case of Hmel tumorigenic the correlation increased to 0.84. With the addition of the fourth attribute, non-tumor condition and Pmel tumorigenic the correlation increased to 0.94 and 0.87, but Hmel tumorigenic decreased to 0.84. With the fifth attribute, Pearson's coefficient increased in all conditions: for non-tumor to 0.95 and for Hmel tumorigenic and Pmel tumorigenic to 0.85 and 0.90. Moreover, we also obtained an improvement with 6 attributes in the tumor conditions to 0.86 and 0.91 for Hmel and Pmel tumorigenic respectively, and in the case of the non-tumor line the value of the previous correlation was maintained. Finally, the models with 7 and 8 attributes show no variation in their results with respect to the 6-attribute model.

#### 1.1.3 H3K4ME3

We obtained a Pearson correlation of 0.77 with the model created using the most importnat attribute to predict H3K4ME2 for the non-tumor condition, correlations of 0.63 and 0.46 were obtained for Hmel and Pmel tumorigenic. With the addition of the second most important attribute, a value of 0.82 is obtained for the non-tumor condition, respectively for 0.72 and 0.61 for Hmel and Pmel tumorigenic. With the addition of the third attribute, we see a drop in the correlation values for the three conditions used, which changes with the addition of the fourth attribute but is still lower than the model that uses only two attributes. Adding the fifth attribute shows an increase in the results with 0.86 for the non-tumor condition, 0.80 and 0.64 with Hmel and Pmel tumorigenic. With the sixth attribute there are only variations in Pmel tumorigenic, with a relative small increase to a correlation of 0.68 is seen. Finally, with the addition of the seventh attribute, the results are maintained except for Pmel tumrigenic where it decreases to 0.67, in the case of the model with 8 attributes, the values are maintained for the model with 7 attributes.

#### 1.1.4 H3K27ME3

The model generated only with the most important mark (H3K23AC) to predict this mark gives low Pearson values with 0.51 for the non-tumor condition, 0.49 for tumorigenic Hmel and 0.44 for tumorigenic Pmel. When generating the model with the two most important descriptors (H3K23AC and H3K36ME2), we obtained an increase in the correlation to 0.72 for the non-tumor condition, 0.71 and 0.68 respectively for Hmel and Pmel tumorigenic, whic indicates a possible relationship of interdependence of H3K23AC and H3K36ME2 with H3K27ME3. When adding the third mark, an abrupt decrease in the results is seen in all conditions, reaching a correlation of 0.06 in Pmel tumorigenic, which decreases even more when adding the fourth mark, reaching 0.03 in the same condition. With the addition of the fifth mark, a great increase in the results is seen, surpassing those obtained in the model generated with the two most important marks, reaching correlations of 0.80 for the non-tumor condition, 0.82 and 0.77 for Hmel and Pmel tumorigenic. With the addition of the sixth attribute, no greater variation is seen compared to the previous model, the values are maintained. By adding the seventh attribute, a new increase in the results is seen with correlations of 0.81, 0.83 and 0.78 for the non-tumor condition, Hmel tumorigenic and Pmel tumorigenic. Finally, with the addition of the eighth attribute, no variation is seen with respect to the previous model.

#### 1.1.5 H3K36ME2

For H3K36ME2, the RF model generated with the most important attribute (H4K8AC) gave Pearson values of 0.75 in the non-tumor condition, 0.75 and 0.68 in tumorigenic Hmel and Pmel. When adding the second most important attribute (H3K36ME1), an increase in the results is observed with correlations of 0.83 for the non-tumor condition, 0.81 and 0.79 with tumorigenic Hmel and Pmel, indicating the exitence of a possible relationship of interdependence with H4K8AC and H3K36ME1 with H3K36ME2. When adding the third and fourth attribute, a decrease is observed in all cell lines, which increases when adding the fifth attribute, reaching correlation values of 0.87 in the non-tumor condition, 0.87 and 0.82 in tumorigenic Hmel and Pmel. Adding the sixth attribute maintains the results in the non-tumor condition, but a slight increase is seen in both tumor lines (tumorigenic Hmel, 0.88 and tumorigenic Pmel, 0.83). With the addition of the seventh attribute, the results are maintained except in Pmel tumorigenic, where a slight increase (0.84) was seen. Finally, when adding attribute 8, a decrease is observed in all cell lines, except in Pmel tumorigenic, where it remained.

#### 1.1.6 H4K20ME2

The model generated with the most important attribute for this mark (H4K20ME3), gives correlation values of 0.71 for the non-tumor condition, 0.67 and 0.65 for tumorigenic Hmel and tumorigenic Pmel. With the addition of the second top attribute (H3K4AC), a slight increase is seen in all conditions, obtaining values of 0.74 for the non-tumor condition, 0.69 and 0.68 respectively for tumorigenic Hmel and tumorigenic Pmel. With the third and fourth attributes, a decrease is seen in all conditions, which increases with the addition of the fifth attribute, obtaining correlation values of 0.76 for the non-tumor condition, 0.71 and 0.69 for tumorigenic Hmel and tumorigenic Pmel. With the sixth attribute, there is only an increase in correlation for tumorigenic Hmel, reaching a value of 0.72. Finally, with the seventh and eighth attributes, no variations are seen with respect to the 6-attribute model.

#### 1.1.7 H4K20ME3

Using only the most important attribute to preditct this mark (H4K20ME2), correlations values are 0.69 for the non-tumor condition, 0.68 and 0.67 for tumorigenic Hmel and tumorigenic Pmel. With the addition of the second best attribute (H3K4AC), we observe an increase in the correlation in all conditions, obtaining 0.71 for the non-tumoral, 0.71 and 0.69 with tumorigenic Hmel and tumorigenic Pmel. With the third attribute correlation decreases in all conditions and increases again when adding the fourth attribute, but is still low compared to the two-attribute model. With the fifth attribute, a new increase is seen, obtaining values of 0.74 for the non-tumor condition, 0.74 and 0.72 for tumorigenic Hmel and tumorigenic Pmel, respectively. With the addition of the sixth attribute only a small increase in tumorigenic Hmel is seen with 0.75.

Finally, it is seen that the results hold with the addition of the seventh and eighth attributes.

##### 1.2 Gene Regulatory Network analysis whole genome

A GRN was created using the whole genome data, which consists of 2248 nodes and 61526 connections. Of the total number of nodes, 64 correspond to transcription factors (TFs) (the Cytoscape session [3] with the generated network is attached as supplementary file 1). Additionally, those target genes that are related to colorectal cancer were identified. The CancerGeneticsWeb [4] and DisGeNet [5] databases were used for this. In this way, 114 genes were obtained, which according to the literature, are associated with colorectal cancer (Table S10).

#### 2 Supplementary Tables Captions

Table S1: Initial test Regression models (All depths). First column indicates the fragment size; Second column the distances adjacent to the central fragment; Next columns indicate the Pearson coefficient correlation for the analyzed Random Forest (RFs).

Table S2: Results obtained when analyzing the 33 available post translational modifications (PTMs) with 35 and 8 attributes. These models were trained using the non-tumorigenic cell line Pmel with 35 attributes (32 PTMs and annotation data) and 8 attributes with greater importance according to Random Forest (RF). The models were tested using the data from the non-tumorigenic cell lines Hmel (Hmel-noTum) and the tumorigenic cell lines Hmel and Pmel (Hmel-Tum and Pmel-Tum). The performance of the models was analyzed using Pearson's correlation coefficient.

Table S3: Values of importance obtained through the analysis with Random Forest (RF) regressor. The first column indicates the post translational modification (PTM) to predict, the following columns indicate the importance values of the attributes used. The importance values are in decreasing order.

Table S4: Test whole genome of 33 available post translational modifications (PTMs) with 35 and 8 attributes. These models were trained using the non-tumorigenic cell line Pmel with 35 attributes (32 PTMs and annotation data) and 8 attributes with greater importance according to Random Forest (RF). The models were tested using the data from the non-tumorigenic cell lines Hmel (Hmel-noTum) and the tumorigenic cell lines Hmel and Pmel (Hmel-Tum and Pmel-Tum). The performance of the models was analyzed using Pearson's correlation coefficient.

Table S5: Models obtained for the different groups identified, using only three shared attributes to predict each mark. The models were trained using the non-tumorigenic cell line Pmel (Pmel-noTum) with the three shared attributes of each group, the models were tested using the data from the non-tumorigenic cell lines Hmel (Hmel-noTum) and the tumorigenic cell lines Hmel and Pmel (Hmel-Tum and Pmel-Tum). The performance of the models was analyzed using Pearson's correlation coefficient.

Table S6: Results obtained when analyzing the 33 post translational modifications (PTMs), increasing the the amount of attributes. The attributes were increased from the one with the highest importance until reaching the eighth. The models were trained using the non-tumorigenic cell line Pmel (Pmel-noTum) with the three shared attributes of each group, the models were tested using the data from the non-tumorigenic cell lines Hmel (Hmel-noTum) and the tumorigenic cell lines Hmel and Pmel (Hmel-Tum and Pmel-Tum). The performance of the models was analyzed using Pearson's correlation coefficient.

Table S7: T-test significance analysis of the results obtained from the comparisons with other methods. The first column indicates the predicted post translational modification (PTM), the following ones indicate the p-values obtained with the comparisons of the different methods, using all 8 and 3 attributes.

Table S8: Comparison with other methods (All PTMs). The first column indicate the PTM predicted. The next columns show the results of the generated models using different amount of attributes. The performance of the models was analyzed using Pearson's correlation coefficient. In bold number indicate the method with statistically significant better results using a t-test (p-value $\leq$ 0.05)

Table S9: Biological process enrichment analysis with Enrichr. Output table obtained by analyzing the target genes of the Gene Regulatory Networks (GRNs) with Enrichr.

Table S10: List of genes associated to colorectal cancer. Target genes obtained with the analysis of GRNs that are associated with colorectal cancer according to CancerGeneticsWeb [4] and DisGeNet [5].
